# Endocytosis-mediated vitellogenin absorption and lipid metabolism in the hindgut-derived pseudoplacenta of the viviparous teleost *Xenotoca eiseni*

**DOI:** 10.1101/2022.02.16.480647

**Authors:** Atsuo Iida, Jumpei Nomura, Junki Yoshida, Takayuki Suzuki, Hayato Yokoi, Eiichi Hondo

**Affiliations:** Laboratory of Animal Morphology, Graduate School of Bioagricultural Sciences, Nagoya University, Tokai National Higher Education and Research System, Nagoya, Aichi, Japan; Laboratory of Marine Life Science and Genetics, Graduate School of Agricultural Science, Tohoku University, Sendai, Miyagi, Japan

## Abstract

Certain viviparous animals possess mechanisms for mother-to-embryo nutrient transport during gestation. *Xenotoca eiseni* is one such viviparous teleost species in which the mother supplies proteins and other components to the offspring developing in the ovary. The embryo possesses trophotenia, a hindgut-derived pseudoplacenta to receive the maternal supplement. However, the molecular mechanisms underlying viviparous non-mammalian animals remain elusive. We conducted this study to investigate the mechanism for nutrient absorption and degradation in trophoenia of *X. eiseni*. The tracer assay indicated that a lipid transfer protein, vitellogenin (Vtg), was absorbed into the epithelial layer cells of trophotaenia. Vtg uptake was significantly suppressed by Pitstop-2, an inhibitor of clathrin-mediated endocytosis. Gene expression analysis indicated that the genes involved in endocytosis-mediated lipolysis and lysosomal cholesterol transport were expressed in trophotaenia. In contrast, plasma membrane transporters expressed in the intestinal tract were not functional in trophotaenia. Our results suggested that endocytosis-mediated lysosomal lipolysis is one of the mechanisms underlying maternal component metabolism. Thus, our study demonstrated how viviparous teleost species have acquired a unique developmental system that is based on the hindgut-derived pseudoplacenta.

## Introduction

For viviparous animals, mother-to-embryo nutrition supply is important for embryonic development and growth in the mother’s body. In vertebrate evolution, more than a hundred viviparous acquisition events are presumed to have occurred in each taxon [1, 2]. Other than mammals, mother-to-embryo nutrient transport has been reported in viviparous species such as bony fishes, cartilaginous fishes, amphibians, and reptiles [3]. The transport machinery varies among species, and organ analogous to the mammalian placenta have been reported in fish, amphibians, and reptiles [4]. In viviparous mammals, embryo-derived placental tissues interact with or fuse with the deciduous membrane of the mother [5]. In contrast, certain non-mammalian viviparous species such as the viviparous teleost belonging to the family Goodeidae possess a unique pseudoplacenta that makes no direct contact with the mother tissue rather absorbs the maternal supplements in the form of secreted factors in the ovarian or uterine luminal fluids [6, 7]. Thus, non-mammalian viviparous species have acquired unique machineries for mother-to-embryo nutrition supply, and their molecular mechanisms are predicted to differ from those of viviparous mammals. However, the molecular mechanisms underlying viviparous non-mammalian animals remain elusive. We conducted this study to investigate the absorption machinery of maternal nutrients in the pseudoplacenta of a viviparous teleost species, *Xenotoca eiseni*.

*Xenotoca eiseni* is a viviparous teleost species belonging to the family Goodeidae, a small freshwater fish distributed in Mexico [8, 9]. *X. eiseni* and other goodeid species are known to possess hindgut-derived pseudoplacenta called trophotaenia. Trophoaeniae are ribbon-like structures elongated from around the anus of the intraovarian embryo [7]. The pseudoplacenta is considered to contribute to the absorption of maternal nutrients during embryonic growth in the ovary [10, 11]. Histological studies have suggested that the absorption machinery involves intracellular vesicle trafficking [12]. However, the nature of the maternal component and molecules responsible for absorption remain elusive.

Vitellogenin (Vtg), an egg yolk nutrient protein, is conserved in nearly all oviparous vertebrate species, including fish, amphibians, reptiles, birds, and monotremes [13, 14]. Since the mid-1900s, Vtg has been investigated as a matrotrophic nutrient for the intraovarian embryo in viviparous teleosts, including in the family Goodeidae [15–17]. In 2019, for the first time, we demonstrated that the yolk protein Vtg is a maternal component, which is incorporated into the epithelial layer cells of trophotaenia using *X. eiseni* via vesicle trafficking [18]. Furthermore, our recent study suggested that vesicle trafficking is driven by receptor-mediated endocytosis (RME). Cubilin-amnionless RME and cathepsin L-dependent proteolysis are active in the epithelial layer cells of trophotaenia [19]. However, the relationship between Vtg and RME has not been demonstrated experimentally. Furthermore, Vtg is a lipid transfer protein carrying phospholipids, triglycerides, and cholesterol [20–22]. Thus, lipids are also possible candidates for maternal nutrient metabolism in trophotaenia. In this study, we investigated the mechanisms of endocytosis, lipolysis, and the vesicle trafficking pathway in *X. eiseni* trophotaenia.

## Methods

### Animal experiments

This study was approved by the Ethics Review Board for Animal Experiments of Nagoya University (A210264-001). A minimal number of live animals were euthanized under anesthesia according to institutional guidelines.

### Fishes

*X. eiseni* was purchased from Meito Suien Co. Ltd. (Nagoya, Japan). The fishes used in this study were maintained in a freshwater circulation system at 27 °C and a 14:10 h light:dark photoperiod cycle.

### Sample collection

Fertile females were examined in the study. The fish samples were anesthetized using tricaine (Nacalai, Kyoto, Japan) on ice prior to the surgical extraction of tissues or embryos. The obtained samples were stored on ice until subsequent experiments. Approximately 10 *X. eiseni* females and their offspring (10–30 embryos per female) were dissected in the study.

### RNA-Sequencing (RNA-Seq)

The transcriptome datasets used in this study were obtained from the DNA Databank of Japan (DDBJ: DRA011209 and DRA011388). Detailed experimental information has been described in our previous publications [19, 23]. To calculate transcript per million values, transcript sequences of *Poecilia reticulata* (National Center for Biotechnology Information [NCBI] Genome, ID: 23338) were used as a reference. *De novo* assembly and mapping to the reference sequence were performed using CLC Genomics Workbench (Filgen Inc., Nagoya, Japan). The coding sequences determined in this study were deposited in DDBJ (Table S1).

### Tracing of Vtg-fluorescein isothiocyanate (FITC)

FITC-conjugated goldfish Vtg protein was prepared according to a previous study [24]. Vtg-FITC was diluted in 100 µg/mL phosphate-buffered saline (PBS), and intra-ovarian embryos were extracted from pregnant female fish at three weeks post-mating. The obtained embryos were incubated in Vtg-FITC solution or PBS for 1 h. In the case of Pitstop-2 exposure, the embryos were pre-incubated in 30 µM Pitstop-2 or the negative control/PBS for 30 min prior to Vtg-FITC treatment. The samples were fixed with 4.0% PFA/PBS, and they were used for paraffin sectioning and immunohistochemistry (IHC).

### Immunohistochemistry

Deparaffinized section samples were permeabilized using 0.5% TritonX-100/PBS at room temperature for 30 min. Endogenous peroxidase was inactivated using 3.0% hydrogen peroxide/PBS for 10 min. The sample was treated with Blocking-One solution (Nacalai, Kyoto, Japan) at room temperature for 1 h. Primary antibodies or antisera were used at 1:500 dilution with Blocking-One solution. The samples were incubated with primary antibody or antiserum at 4 °C for 16 h. Secondary antibodies were used at 1:500 dilution in 0.1% Tween-20/PBS. The samples were then treated with a secondary antibody solution at 4 °C for 2 h. 3,3’-diaminobenzidine tetrahydrochloride (DAB) color development was performed using the DAB Peroxidase Substrate Kit, ImmPACT (Vector Laboratories, Inc., Burlingame, CA, USA), as per manufacturer instructions. In case of the fluorescent immunohistochemistry (FITC), the nucleus was stained using DAPI (Sigma, St. Louis, MO, USA), and filamentous actin (F-actin) was stained using Alexa Fluor® 546 Phalloidin (Thermo Fisher Scientific, Waltham, MA, USA). Microscopic observations were performed using an Olympus BX53 microscope, and images were photographed using a DP25 digital camera (Olympus, Shinjuku, Japan), BZ-X800 microscope (Keyence, Osaka, Japan) or a DM5000 B microscope (Leica Microsystems). The antibodies used in this study are listed in Table S2.

### Labeling acidic organelles

Immediately after extraction from the pregnant female at three weeks post-mating, live embryos were incubated in PBS with a 1:1000 dilution of LysoTracker® Red (Thermo Fisher Scientific, Waltham, MA, USA) at 20 °C. for 1 h. The samples were fixed with 4.0% PFA/PBS and stained with DAPI. Microscopic observations were performed using a DM5000 B microscope (Leica Microsystems).

### Domain search

Amino acid sequences obtained from the *de novo* assembly of *scarb1*, *lipa*, *npc1*, *npc2*, *npc1l1*, *cd36*, *slc27a1*, and *slc27a4* were surveyed using PROSITE (https://prosite.expasy.org/) [25], TMHMM-2.0 (https://services.healthtech.dtu.dk/service.php?TMHMM-2.0) [26], and SignalP-5.0 (https://services.healthtech.dtu.dk/service.php?SignalP-5.0) [27] to predict functional domains, transmembrane regions, and signal peptides.

### Phylogenetic analysis

Amino acid sequences for *scarb1*, *lipa*, *npc1*, *npc2*, *npc1l1*, *cd36*, *slc27a1*, and *slc27a4* of *Homo sapiens* (human), *Mus musculus* (house mouse), *Oryzias latipes* (medakafish), *P. reticulata* (guppy), and *X. eiseni* were collected from the NCBI protein database (https://www.ncbi.nlm.nih.gov/ protein/). Phylogenetic trees were constructed using the neighbor-joining method with 500 bootstrap replicates in the MEGAX (version 10.1.8) software (https://www.megasoftware.net/) [28, 29]. The amino acid sequences used in the phylogenetic analysis are listed in Table S3.

### Reverse transcription-polymerase chain reaction (RT-PCR)

The total RNA content was extracted from adult and embryonic tissues using the RNeasy Plus Mini kit (Qiagen), and it was reverse-transcribed using SuperScript IV reverse transcriptase (Thermo Fisher Scientific). PCR was performed using KOD-FX-Neo (Toyobo, Osaka, Japan) under the following conditions: 94 °C for 100 s, followed by 32 cycles of 94 °C for 20 s, 60 °C for 10 s, 72 °C, and 72 °C for 20 s. The primer sequences are listed in Table S4.

### Quantitative reverse transcription-polymerase chain reaction (qPCR)

qPCR was performed using the Roche LightCycler 96 system (Roche, Mannheim, Germany) with Thunderbird SYBR qPCR Mix (Toyobo, Osaka, Japan) under the following conditions: 40 cycles of 60 s at 95 °C, 30 s at 55 °C, and 30 s at 72 °C. The relative expression level was expressed as the reciprocal of ΔCt. After normalization using Ct for b-actin, the relative expression values for trophotaenia and adult intestines were compared. The primer sequences are listed in Table S5.

## Results

### Clathrin-dependent Vtg uptake

In this study, we used the approximately third week embryo after fertilization, which was confirmed via macroscopic observation. Trophotenia was well developed at this stage, and its absorption structure comprised a single layer of epithelial cells (Fig. 1A). Endocytosis-mediated intracellular vesicles in cells were visualized by immunohistochemistry against cubilin, an endocytic receptor expressed in trophotaenia (Fig. 1B) [19]. To trace Vtg absorption, embryos were extracted from pregnant female fish and incubated in PBS containing Vtg-FITC for 1 h. In the case of clathrin inhibition, embryos were exposed to Pitstop-2 for 30 min prior to Vtg addition (Fig. 1C). IHC against FITC revealed the distribution of conjugated protein in trophotaenia (Fig 1D). Signal accumulation was observed in the epithelial layer of trophotaenia, which showed a similar pattern to that of IHC against cubilin (Fig. 1E). As a functional factor for the endocytosis of Vtg, we focused on the major vesicle-coating protein clathrin, which consists of heavy- and light-chain subunits. In *X. eiseni*, clathrin genes *clta* and *cltc* were expressed in trophotaenia (Fig. 1F). *Clta* and *cltc* were detected in the adult intestine as these genes contributed to intestinal absorption. Muscle samples were used as a negative control for endocytosis-negative tissues. The full-length amino acid sequences for clta and cltc of *X. eiseni* were determined and compared with those of the clathrin proteins of *H. sapiens*. Particularly, the terminal domain was highly conserved (Fig. 1G). *X. eiseni* clathrin proteins were closest to those of *P. reticulata* in the four vertebrate species *H. sapiens*, *M. musculus*, *O. latipes*, and *P. reticulata* (Fig. S1B). For functional analyses, we used Pitstop-2, which is a cell-permeable clathrin-mediated endocytosis inhibitor [30]. Pitsop-2 bonded to the amino acid motif in the terminal domain of clathrin heavy chain (Fig. 1G). The purchased Pitstop-2 compound and negative control analog were used for *ex vivo* exposure analysis of *X. eiseni* embryos (Fig. 1H). Anti-FITC signals in the epithelial layer of trophotaenia were grossly reduced under Pitstop-2 treatment (Fig. 1I), and the negative control exposure did not interfere with the signal distribution. The tissue structure of trophotaenia exhibited no morphological abnormalities due to drug treatment. Lysotracker is an acidic organelle- staining dye that visualizes lysosomal vesicles in the epithelial layer cells of trophotaenia. Part of the red fluorescent signals indicated that the vesicles merged with the green fluorescence, indicating Vtg-FITC. Pitstop-2 exposure reduced the fluorescent signals for both lysotracker and FITC in trophotaenia (Fig 1J, Fig. S1C).

**Figure 1.**
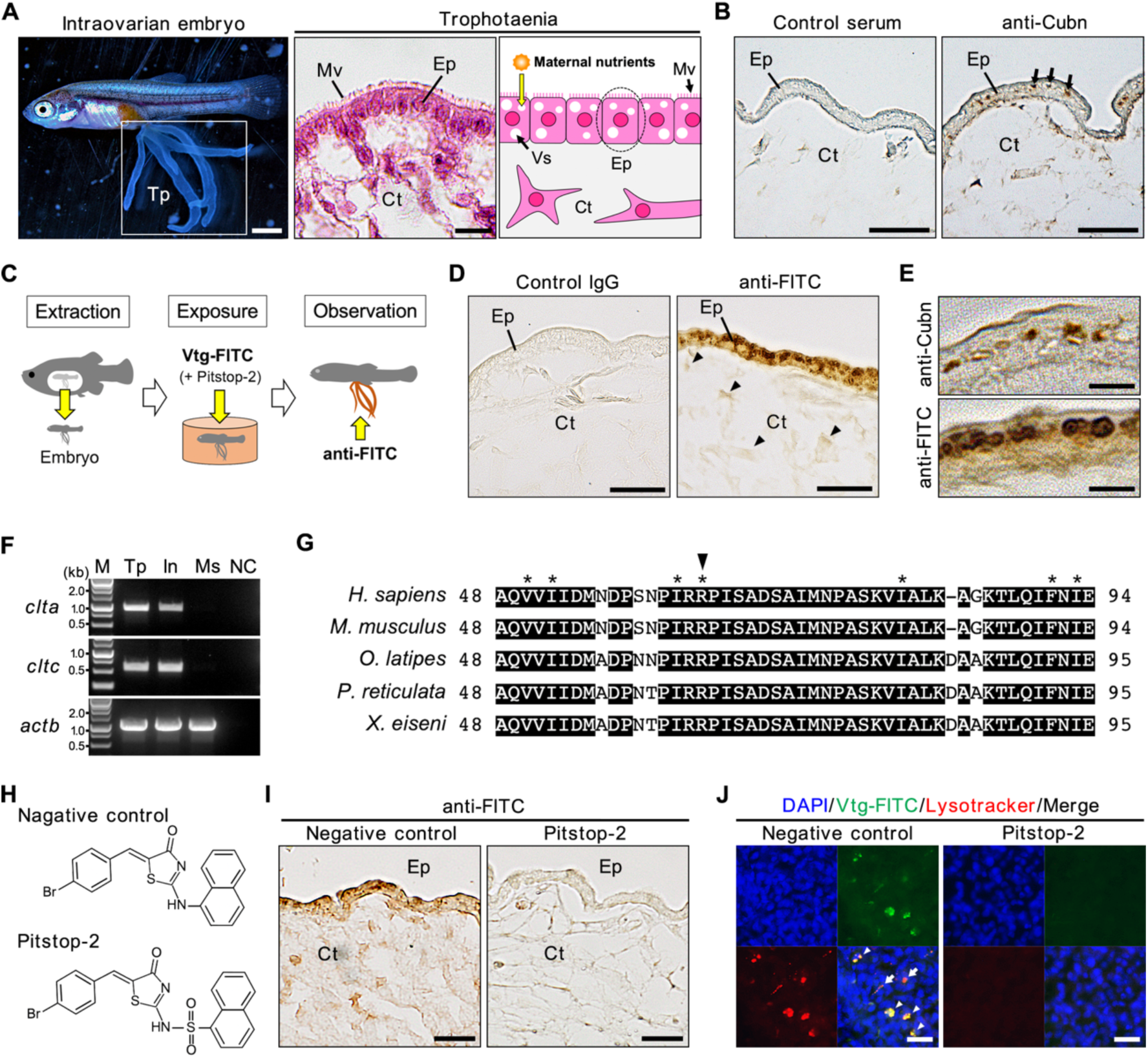
Detection of clathrin-dependent vitellogenin (Vtg) uptake in trophotaenia. **A**. Morphological observation of trophotaenia in the intraovarian embryo of *X. eiseni* at the third week post-fertilization. The section image indicates the tissue structure of trophotaenia. The illustration is a model for maternal nutrient absorption via vesicle trafficking. Scale bar, 1 mm (left) and 10 µm (right). **B**. The sectional image of trophotaenia for immunohistochemistry (IHC) against cubilin. The anti-Cubn signals in the epithelial layer cells indicate intracellular vesicles (arrows). Scale bar, 20 µm. **C**. Graphical abstract for the experimental procedure of Vtg tracing and drug exposure. **D**. The sectional image of trophotaenia for IHC using fluorescein isothiocyanate (FITC) antibody (anti-FITC) or control IgG. The strong anti-FITC signals in the epithelial layer cells indicate Vtg-FITC absorption. The arrowheads indicate weaker signals in the connective tissue cells. Scale bar, 20 µm. **E**. Comparison of the intracellular signals against cubilin and FITC. The strong signals in the epithelial layer cells indicate the intracellular vesicles. Scale bar, 5 µm. **F**. Electrophoresis for RT-PCR to amplify the clathrin genes (*clta* and *cltc*) from the trophotaenia (Tp), adult intestine (In), or adult muscle (Ms) of *X. eiseni*. b-actin (*actb*) was used as the positive control. M, size marker. NC, negative control (no template DNA). **G**. Comparison of amino acid sequences for the N-terminal region of clathrin heavy chain in five vertebrate species, *Homo sapiens*, *Mus musculus*, *Oryzias latipes*, *Poecillia reticulata*, and *Xenotoca eiseni*. The arrowhead indicates the core arginine to bind the Pitstop-2 molecule. Asterisks indicate the amino acids of the binding motif. **H**. Molecular structure for the negative control analog and Pitstop-2. **I**. The sectional image of trophotaenia for IHC using the FITC antibody under negative control or Pitstop-2 exposure. The anti-FITC signals were decreased following Pitstop-2 treatment. Scale bar, 20 µm. **J**. Fluorescence observation for the Lysotracker-stained acidic organelle and absorbed Vtg-FITC in the epithelial layer of trophotaenia. DAPI staining indicates the cell nucleus. Arrowheads indicate the lysosomal vesicles, including strong and few fluorescence for Vtg-FITC. Scale bar, 10 µm. Ct, connective tissue. Ep, epithelial layer cell. Mv, microvilli. Vs, intracellular vesicle.

### Endocytosis-mediated intracellular lipolysis

Endocytosis-mediated intracellular lipolysis is a lipid trafficking pathway driven by endocytosis receptors, digestion enzymes, and lipid transport mediators (Fig. 2A). From the open transcriptome data, candidate genes for endocytosis-mediated lipid metabolism expressed in trophotaenia were isolated and their expression levels were determined (Fig. 2B, Table S6). Scavenger receptor class b member 1 (scarb1) is a scavenger receptor for various lipoproteins, similar to cubilin, as previously described (Iida et al., 2021). Lysosomal acid lipase (lipa) is a hydrolase for triglycerides or cholesterol esters that is optimal under acidic conditions, such as the environment in lysosomal vesicles. Niemann–Pick types c1 and -2 transport cholesterol from the lysosomal lumen to cytosol. Amino acid sequences of the gene products of *X. eiseni* were closest to those of *P. reticulata* in four vertebrate species, *H. sapiens*, *M. musculus*, *O. latipes*, and *P. reticulata* (Fig. S2A-D, Table S3). RT-PCR indicated that all genes were expressed in the trophotaenia of intraovarian embryos and adult intestines (Fig. 2C). In contrast, the expression of endocytic genes was not detected in adult muscles. The functional domains of these proteins were conserved between *H. sapiens* and *X. eiseni,* which included cholesterol binding, lysosomal targeting, or lipase activity motifs (Fig. 2D-G, Fig. S2E-G). The fluorescent immunohistochemistry indicated that the lipa protein is distributed in the extranuclear cytosol in the epithelial layer cells (Fig. 2H, Fig. S2H).

**Figure 2.**
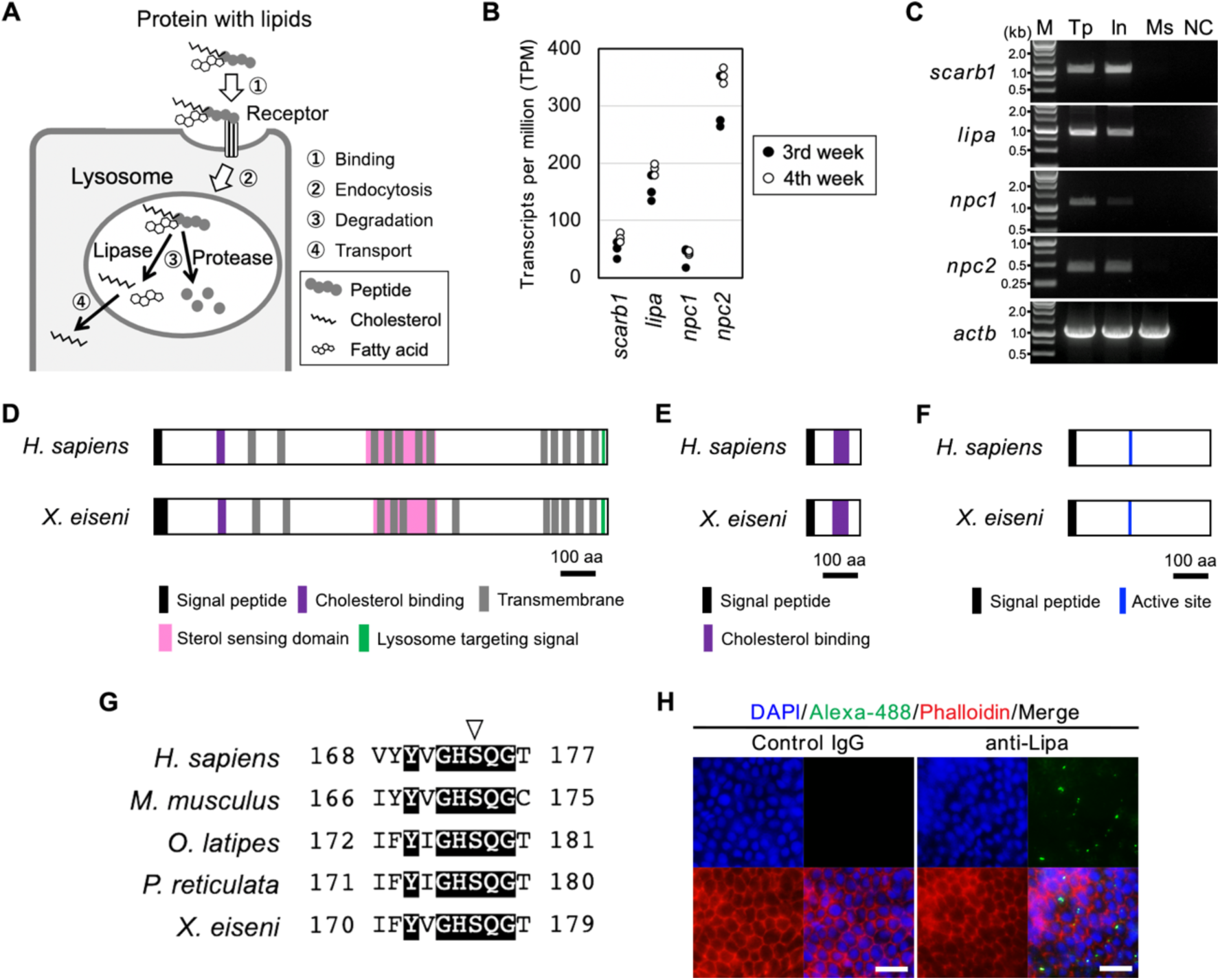
Validation of endocytosis-mediated lipolysis in trophotaenia. **A**. A working hypothesis for endocytosis-mediated lipoprotein absorption and degradation. **B**. Comparison of the expression for endocytosis-related genes (*scarb1*, *lipa*, *npc1*, and *npc2*) in trophotaenia. The expression values are transcripts per million obtained from the published RNA-Seq dataset of 3^rd^- or 4^th^ week trophotaenia. **C**. Electrophoresis for RT-PCR to amplify the endocytic genes in trophotaenia (Tp), adult intestine (In), or adult muscle (Ms) of *X. eiseni*. b-actin (*actb*) was used as the positive control. M, size marker. NC, negative control (no template DNA). **D-F**. Comparison of the structures for npc1 (D), npc2 (E), and lipa (F) protein between *Homo sapiens* and *Xenotoca eiseni*. The functions for cellular distribution, lipid binding, or lipolytic activity were predicted to be conserved between the species. **G**. Comparison of the amino acid sequences for the lipase domain of lipa between *Homo sapiens*, *Mus musculus*, *Oryzias latipes*, *Poecillia reticulata* and *Xennotoca eiseni*. The white arrowhead indicates a serine residue for the enzyme active center. **H**. The Fluorescent immunohistochemistry (FIHC) against lipa. The anti-Lipa signals (Alexa-488) in the extranuclear cytosol. The phalloidin indicates filamentous actin (F-actin). Scale bar, 20 µm.

### Plasma membrane transport

Plasma membrane transport is another pathway for cytosolic lipid absorption (Fig. 3A). The candidate genes for this pathway were isolated, and the expression levels in trophotaenia were estimated from the open transcriptome data. The membrane protein niemann–pick type c1-like 1 (npc1l1) is a cholesterol transporter, whereas platelet glycoprotein 4 (cd36), solute carrier family 27 member 1 (slc27a1), and solute carrier family 27 member 4 (slc27a4) are fatty acid transporters. Amino acid sequences of the gene products of *X. eiseni* were closest to those of *P. reticulata* in four vertebrate species, *H. sapiens*, *M. musculus*, *O. latipes*, and *P. reticulata* (Fig. S3A-D). RNA-seq indicated that all the above-mentioned genes, except *slc27a4*, were only weakly expressed in trophotaenia of the intraovarian embryo (Fig. 3B). *npc1l1* and *cd36* genes were not detected in trophotaenia using RT-PCR (Fig. 3C). qPCR indicated that the expression of each gene in trophotaenia was significantly lower than that in the adult intestine (Fig. 3D). Although the expression of *slc27a1* and *slc27a4* was detected in trophotaenia via RT-PCR, some of the functional domains of gene products were not conserved between *H. sapiens* and *X. eiseni*. Computational prediction indicated that *X. eiseni* slc27a1 lacked the lipocalin domain, which is required for lipid binding, and it possessed an additional transmembrane region as compared to the *H. sapiens* protein (Fig. 3E, Fig. S3E and G). *X. eiseni* slc27a4 had no transmembrane region; thus, it was considered to be a secreted protein (Fig. 3F, Fig. S3F and H). The results indicated that the lipid transport activity driven by the four proteins was incomplete in trophotaenia owing to the low gene expression or mutations in their functional domains.

**Figure 3.**
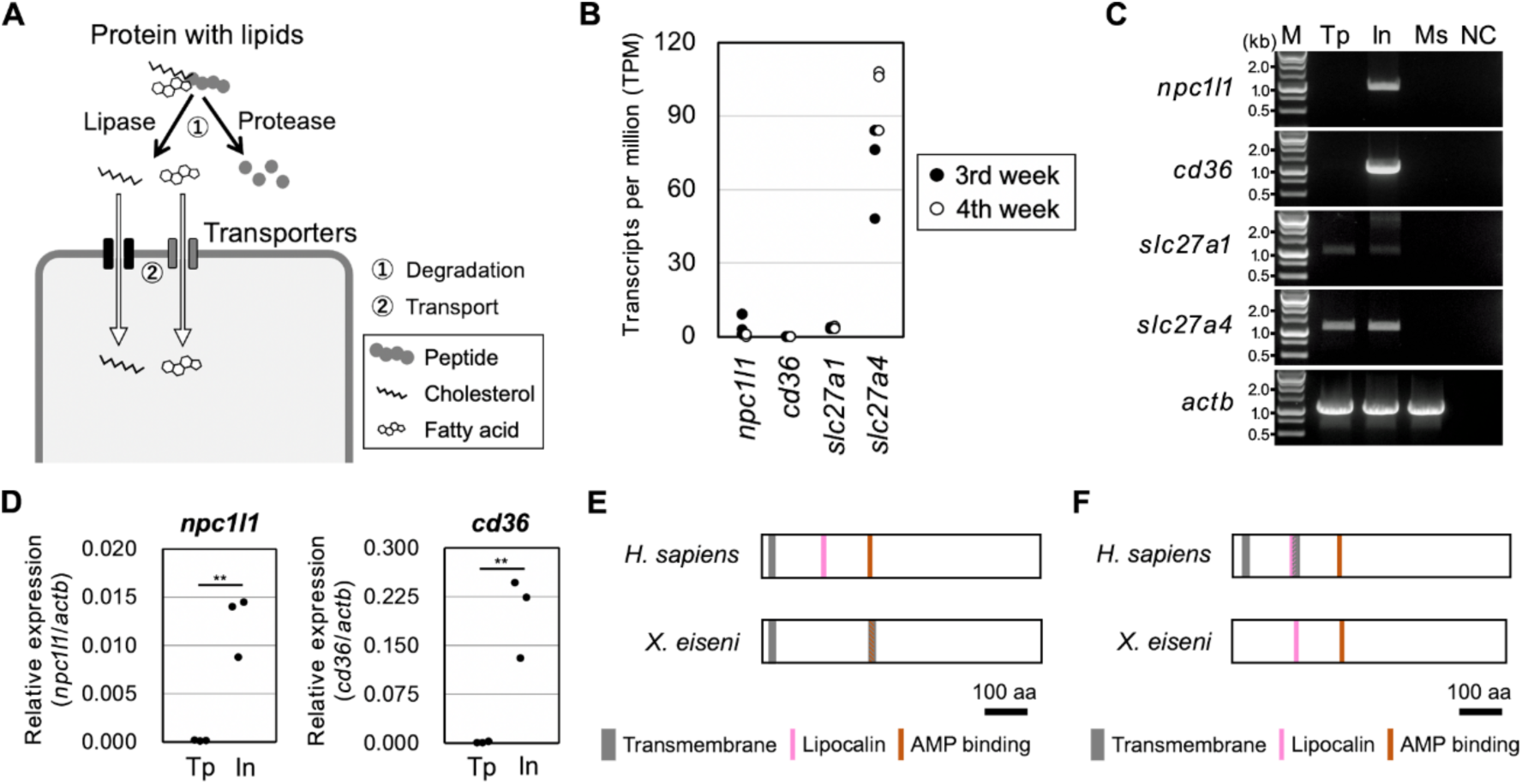
Validation of transporter-mediated lipids uptake in trophotaenia. **A**. A working hypothesis for membrane transporter-mediated lipids transport into the cytosol. **B**. Comparison of the expression of lipid transporter genes (*npc1l1*, *cd36*, *slc27a1*, and *slc27a4*) in trophotaenia. The expression values are transcripts per million obtained from the published RNA-Seq dataset of 3^rd^- or 4^th^ week trophotaenia. **C**. Electrophoresis for RT-PCR to amplify lipid transporter genes in the trophotaenia (Tp), adult intestine (In), or adult muscle (Ms) of *X. eiseni*. b-actin (*actb*) was used as the positive control. M, size marker. NC, negative control (no template DNA). **D**. Comparison of the relative expression values of *npc1l1* or *cd36* between the trophotaenia (Tp) and adult intestine (In). The threshold cycles (Ct values) were used as the gene expression values, and values for the target genes in each tissue were normalized using that for b-actin (*actb*). Student’s *t*-test was used for statistical analyses. \*\**p* < 0.01. **E-F**. Comparison of the structure for slc27a1 (E) and slc27a4 (F) proteins between *Homo sapiens* and *Xenotoca eiseni*. *X. eiseni* slc27a1 lacks the lipocalin domain and possesses an additional transmembrane domain overlapped to the AMP binding motif. *X. eiseni* slc27a4 possessed no transmembrane regions while two distinct hydrophobic domains were observed in the human protein.

## Discussion

This study indicated that *X. eiseni* trophotaenia was involved in clathrin- dependent Vtg uptake and vesicle trafficking. Molecular genetics suggested that the adult intestine possesses both endocytosis-driven protein absorption and membrane transporter-mediated lipid uptake; however, trophotaenia might lack the membrane transport pathway (Fig. 4).

**Figure 4.**
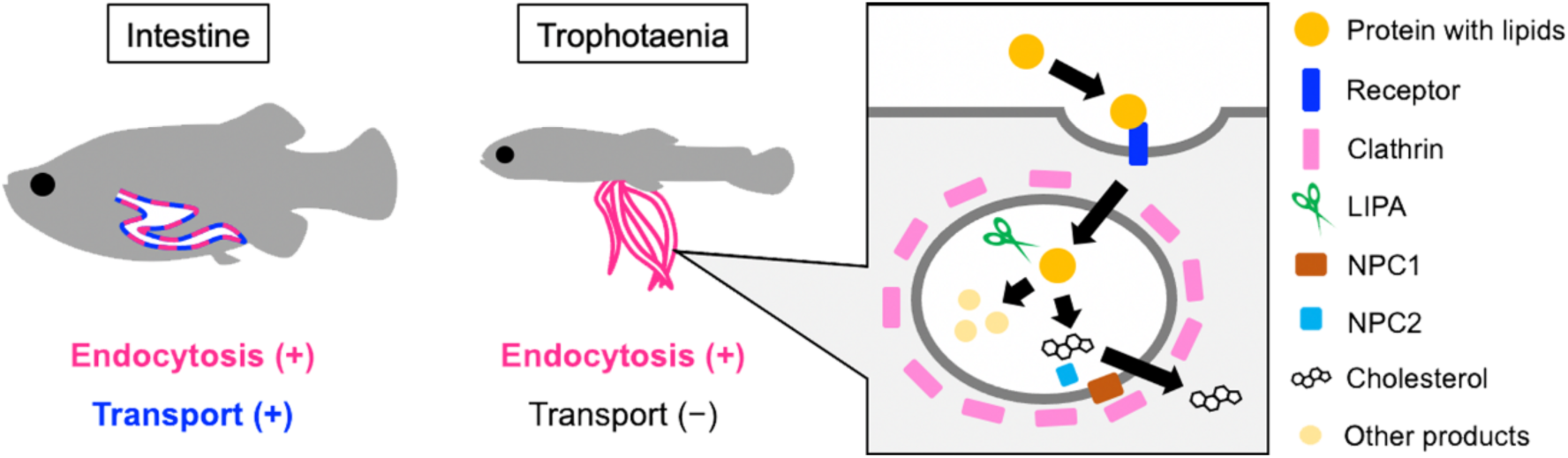
Mechanism of lipid absorption in trophotaenia. Model for lipoprotein absorption and degradation in the adult intestine and embryonic trophotaenia. The molecular pathway based on endocytosis was predicted using genetic and functional assays. These include maternal nutrients, endocytic receptors, vesicle- coating proteins, lipase, and cholesterol mediators.

The absorption of Vtg into trophotaenia was validated using *ex ovo* culture and IHC. This is an improved method from our previous study, based on *in vivo* injection [18]. The Pitstop-2 compound is a well-used compound for inhibition of clathrin- mediated endocytosis [30]. However, the risk of cellular toxicity in non-target tissues cannot be excluded in *in vivo* assays. Considering this concern, quick exposure based on *ex ovo* culture was a suitable method for this study. Pitstop-2 treatment significantly suppressed Vtg absorption without apparent physiological abnormalities, suggesting that Vtg incorporation into the epithelial layer cells of trophotaenia is driven by the clathrin-mediated endocytic pathway. This is the first experimental verification of the driving force of mother-to-embryo nutrient supply in viviparous teleost species. However, a previous study has indicated concerns about Pitsop-2 specificity for clathrin-dependent endocytosis [31, 32]. Unfortunately, our study excluded this possibility. Thus, the conclusion about the factors responsible for this endocytic pathway should be carefully evaluated via further analysis, including genetic manipulation.

Caveolin (cav) and flotillin (flot), major coated vesicle proteins involved in clathrin-independent endocytosis, are possible candidates for this pathway [33–35]. RNA-Seq indicated that *X. eiseni cav2* and *cav3* were not expressed in trophotaenia. In contrast, the expression of *X. eiseni flot1b* and *flot2a* was detected using RNA-Seq (Fig. S4). However, exposure to Pitstop-2 mostly blocked Vtg incorporation into the epidermal layer cells of trophotaenia. A previous study indicated that Pitstop-2 is involved in the positive regulation of vesicles defined by flotillin-2 [36]. Therefore, according to our results, flotillins do not possibly contribute to Vtg intake. Some studies have indicated that glycosylphosphatidylinositol-linked proteins or cholera toxin B subunits are targets for flotillin-mediated endocytosis [34, 37]. We did not exclude the possibility that the flotillin genes expressed in trophotaenia could be involved in the incorporation of co-expressed membrane proteins or maternal components, except Vtg, in the ovarian lumen.

We have previously reported that trophotaenia in intraovarian embryos possesses cathepsin L-dependent protease activity [19]. This activity may be related to the degradation of maternal proteins, including Vtg. However, Vtg is a lipid transport protein; thus, lipolysis and lipid trafficking are also required in trophotaenia. Our molecular genetic analyses suggested that lipa, a lysosomal lipase that hydrolyzes cholesteryl esters and triglycerides and the only known enzyme active at the acidic pH in lysosomes, is active in trophotaenia [38, 39]. The expression of *npc-1* and *-2* suggested that cholesterol transport from the lysosomal lumen to cytosol is active in trophotenia cells [40, 41]. This evidence supported that the epithelial layer cells of trophotaenia possessed both proteolytic and lipolytic activities for lipoproteins absorbed via the RME involved in cubilin, scarb1, or other lipoprotein receptors [42–44].

Trophotaenia is partially derived from the hindgut and exhibits the structural and functional characteristics of intestinal absorptive cells [12, 45]. Our previous study indicated that endocytosis-mediated proteolysis is a common mechanism between intestinal absorbed cells and epithelial layer cells in trophotaenia [18]. In this study, we proposed that endocytosis-mediated lipolysis was a common trait among the absorptive tissues. In contrast, trophotaenia cells might lack the lipid absorption machinery across the plasma membrane via cholesterol or fatty acid transporters while the mechanism was activated in intestinal cells. We hypothesized that the pregnant ovarian lumen contained few digestive enzymes for proteolysis, lipolysis, or glycolysis because it avoids the autolysis of embryos. The small nutrition molecules could be provided from ovarian tissues via direct secretion and not via extracellular digestion in the ovarian lumen; thus, this does not completely explain why trophotenia cells lack plasma membrane transport. On the other hand, our study revealed that endocytosis driven by lysosomal vesicles is a potent pathway for absorption of mother-derived lipoproteins, including Vtg. This finding provides new insights into the viviparity system of goodeid species, which differs from mammals or other viviparous vertebrates.

## Acknowledgments

We thank Yutaka Hattori and Keyence Corporation for complimentary rent to the digital microscope. This work was supported by the Nakatsuji Foresight Foundation, Daiko Foundation.

## Data accessibility

The data that support the findings have been provided with the manuscript.

## Supplemental Information

**Figure S1.**
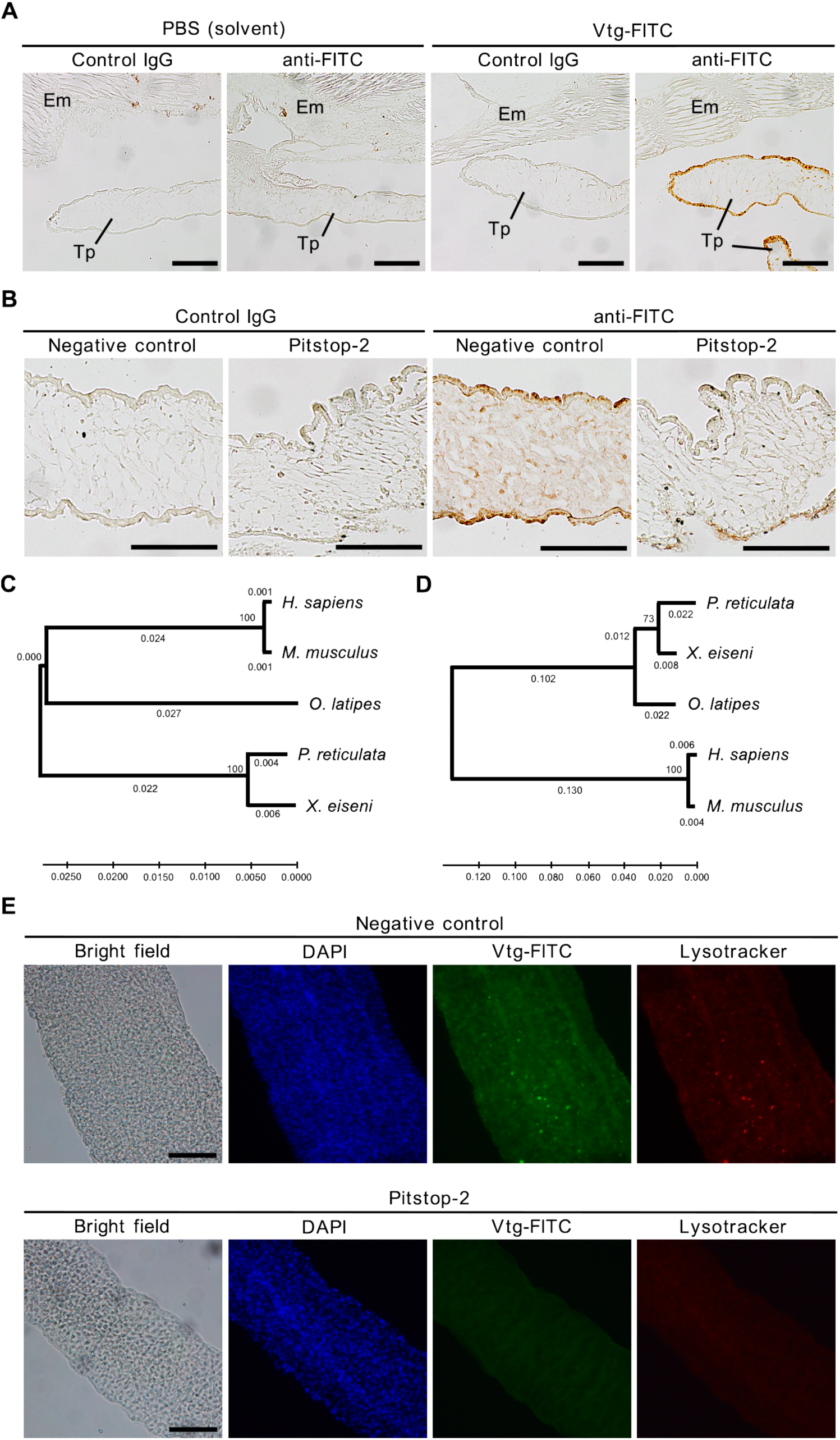
Tracing and inhibition assays related to. Figure 1 **A**. Low-expansion images for Vitellogenin-fluorescein isothiocyanate (Vtg-FITC) tracing in trophotenia. In the solvent treatment, no signal was detected via immunohistochemistry (IHC). In Vtg-FITC exposure, strong signals were detected in the epithelial layer of trophotaenia using the FITC antibody via IHC. The anti-FITC signals were specific to trophotaenia but not to the embryo body. Em, embryo body. Tp, trophotaenia. Scale bar, 100 µm. **B**. Low-expansion images for Vtg-FITC tracing in trophotenia under Pitstop-2 exposure. Scale bar, 100 µm. **C**-**D**. Phylogenetic tree showing the relationship between amino acid sequences of clta (C) and cltc (D) in five vertebrate species (*Xenotoca eiseni*, *Poecilia reticulata*, *Oryzias latipes*, *Homo sapiens*, and *Mus musculus*). The sequences of *X. eiseni* (Cyprinodontiformes: Goodeidae) identified in the present study were close to the sequences of orthologs in *P. reticulata* (Cyprinodontiformes: Poeciliidae). The amino acid sequences used in these calculations are listed in Table S3. **D**. Low expansion images for Vtg-FITC tracing in trophotaenia with acidic organelle labelling using Lysotracker. This is shown in Fig. 1J. Scale bar, 100 µm.

**Figure S2.**
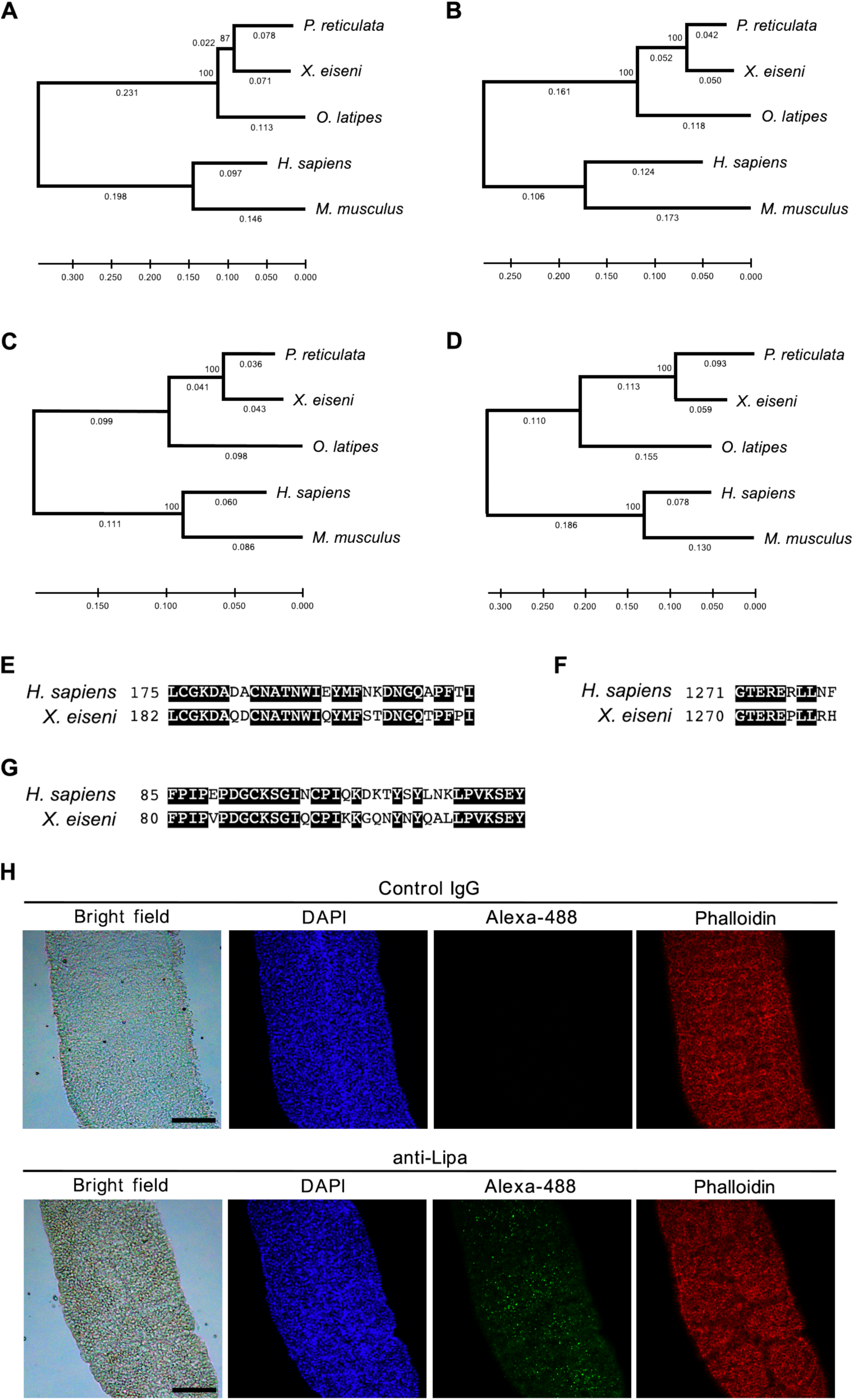
Endocytic factor proteins of *X eiseni* related to. Figure 2**. A**-**D**. Phylogenetic tree indicating the relationship among the amino acid sequences of scarb1 (A), lipa (B), npc1 (C), and npc2 (D) in five vertebrate species (*Xenotoca eiseni*, *Poecilia reticulata*, *Oryzias latipes*, *Homo sapiens*, and *Mus musculus*). All sequences of *X. eiseni* (Cyprinodontiformes: Goodeidae) identified in the present study were close to the sequences of orthologs in *P. reticulata* (Cyprinodontiformes: Poeciliidae). Accession numbers for the amino acid sequences used in the calculation are listed in Table S3. **E**-**F**. Comparison of the amino acid sequences for the cholesterol binding motif (E) and lysosomal targeting motif (F) of npc1 between *Homo sapiens* and *Xennotoca eiseni*. **G**. Comparison of the amino acid sequences for the cholesterol binding motif of npc2 between *Homo sapiens* and *Xennotoca eiseni*. **H**. Low expansion images for fluorescent immunohistochemistry against lipa in trophotaenia with filamentous actin staining using phalloidin. This is shown in Fig. 2H. Scale bar, 100 µm.

**Figure S3.**
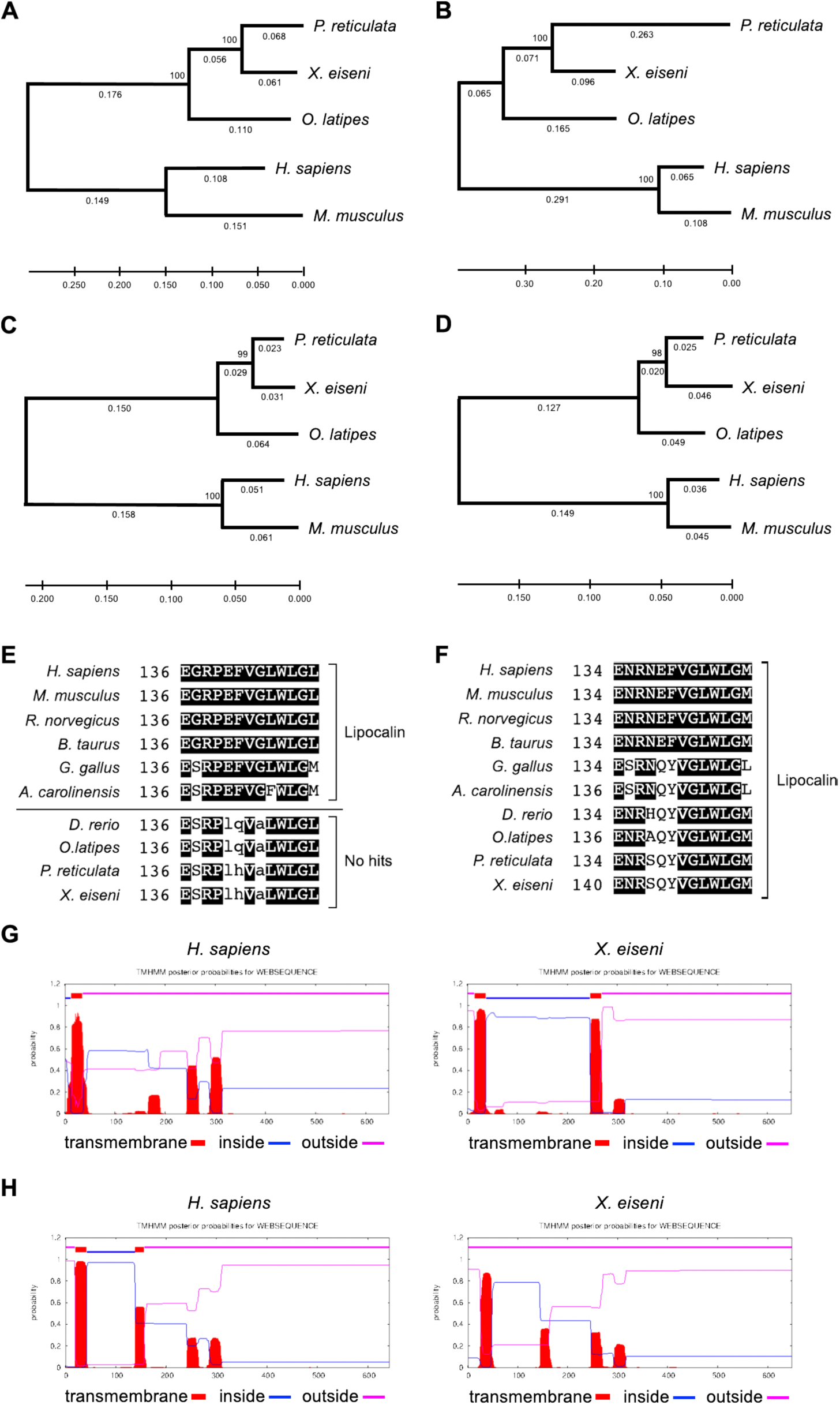
Lipid transporters of *X eiseni* related to. Figure 3 **A-D.** Phylogenetic tree indicating the relationship among the amino acid sequences of npclll (A), cd36 (B), slc27a1 (C), and slc27a4 (D) in five vertebrate species *(Xenotoca eiseni*, *Poecilia reticulata*, *Oryzias latipes*, *Homo sapiens*, and *Mus musculus*). All sequences of *X. eiseni* (Cyprinodontiformes: Goodeidae) identified in the present study were found to be close to the sequences of orthologs in *P. reticulata* (Cyprinodontiformes: Poeciliidae). The amino acid sequences used in the calculation are listed in Table SX. **E**-**F**. Comparison of the amino acid sequences for the lipocalin domain of slc27a1 (E) or slc27a4 (F) in ten vertebrate species, *Homo sapiens*, *Mus musculus*, *Rattus norvegicus*, *Bos taurus*, *Gallus gallus*, *Anolis carolinensis*, *Danio rerio*, *Oryzias latipes*, *Poecillia reticulata*, and *Xenotoca eiseni*. The lowercase letters indicate mismatch amino acids to the lipocalin consensus pattern, [DENG]-{A}- [DENQGSTARK]-x(0,2)-[DENQARK]-[LIVFY]-{CP}-G-{C}-W-[FYWLRH]-{D}- [LIVMTA]. **G-H**. Images for the transmembrane prediction calculated in slc27a1 (G) and slc27a4 (H) via TMHMM-2.0. The program concluded that both proteins exhibited different topologies in the cellular distributions between *H. sapiens* and *X. eiseni*.

**Figure S4.**
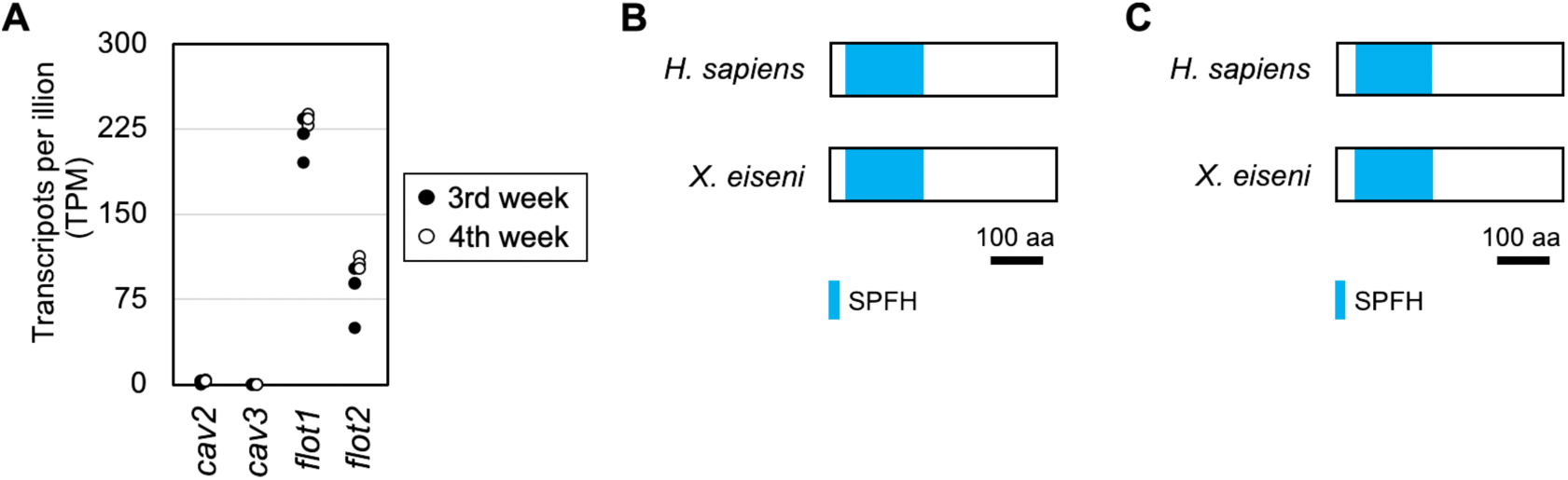
Clathrin-independent endocytosis proteins of *X. eiseni*. **A**. Comparison of the relative expression values of *cav2*, *cav3*, *flot1*, or *flot2* in the trophotaenia of intraovarian embryo. The expression values are transcripts per million obtained from the published RNA-Seq dataset of 3^rd^- or 4^th^-week trophotaenia. **B**-**C**, Comparison of the structures of flot1 (B) or flot2 (C) protein between *Homo sapiens* and *Xenotoca eiseni*. SPFH, stomatin prohibitin flotillin HflK/C.

**Table S1.** The target genes for the nutrient absorption in Xenotoca eiseni

| Name | Symbol | Accession No. | Function |
| --- | --- | --- | --- |
| clathrin heavy chain | clta | LC595292.1 | Intracellular vesicle coating |
| clathrin light chain | cltc | LC595293.1 | Intracellular vesicle coating |
| Scavenger Receptor Class B Member 1 | scarb1 | LC677147.1 | Receptor for high-density lipoprotein |
| lipase A, lysosomal acid type | lipa | LC677148.1 | Hydrolase of cholesteryl esters and triglycerides |
| Niemann–Pick type C1 | npc1 | LC677149.1 | Mediator for intracellular cholesterol trafficking |
| Niemann–Pick type C2 | npc2 | LC677150.1 | Mediator for intracellular cholesterol trafficking |
| NPC1-like 1 | npc1l1 | LC677151.1 | Transporter for cholesterol |
| platelet glycoprotein 4 | cd36 | LC677152.1 | Transporter for fatty acid |
| solute carrier family 27 member 1 | slc27a1 | LC677153.1 | Transporter for long-chain fatty acid |
| solute carrier family 27 member 4 | slc27a4 | LC677154.1 | Transporter for long-chain fatty acid |

**Table S2.** The antibodies used in this study.

| Reagent or Resource | Source | Identifier |
| --- | --- | --- |
| CUBN Polyclonal Antibody | Invitrogen | PA5-83684 |
| Rabbit IgG Isotype Control (Control IgG) | Invitrogen | 02-6102 |
| Goat Anti-Rabbit IgG Antibody (H+L), Peroxidase | Vector | PI-1000 |
| Rabbit polyclonal anti-FITC | Thermo Fisher Scientific | 71-1900 |
| Anti LIPA antibody | Proteintech | 12956-1-AP |
| Donkey polyclonal anti-rabbit IgG (Alexa Fluor® 594 conjugated) | Thermo Fisher Scientific | A21207 |

**Table S3.**
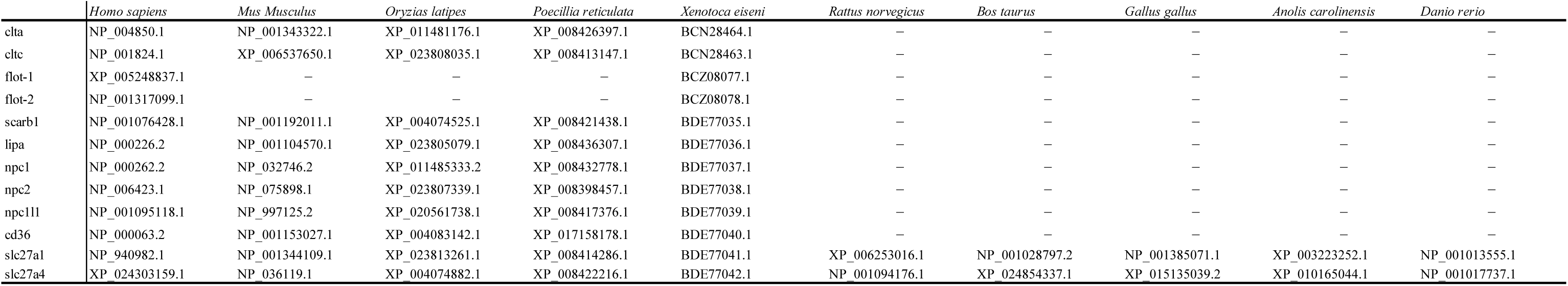
The proteins unsed for phlogenetic analysis.

**Table S4.**
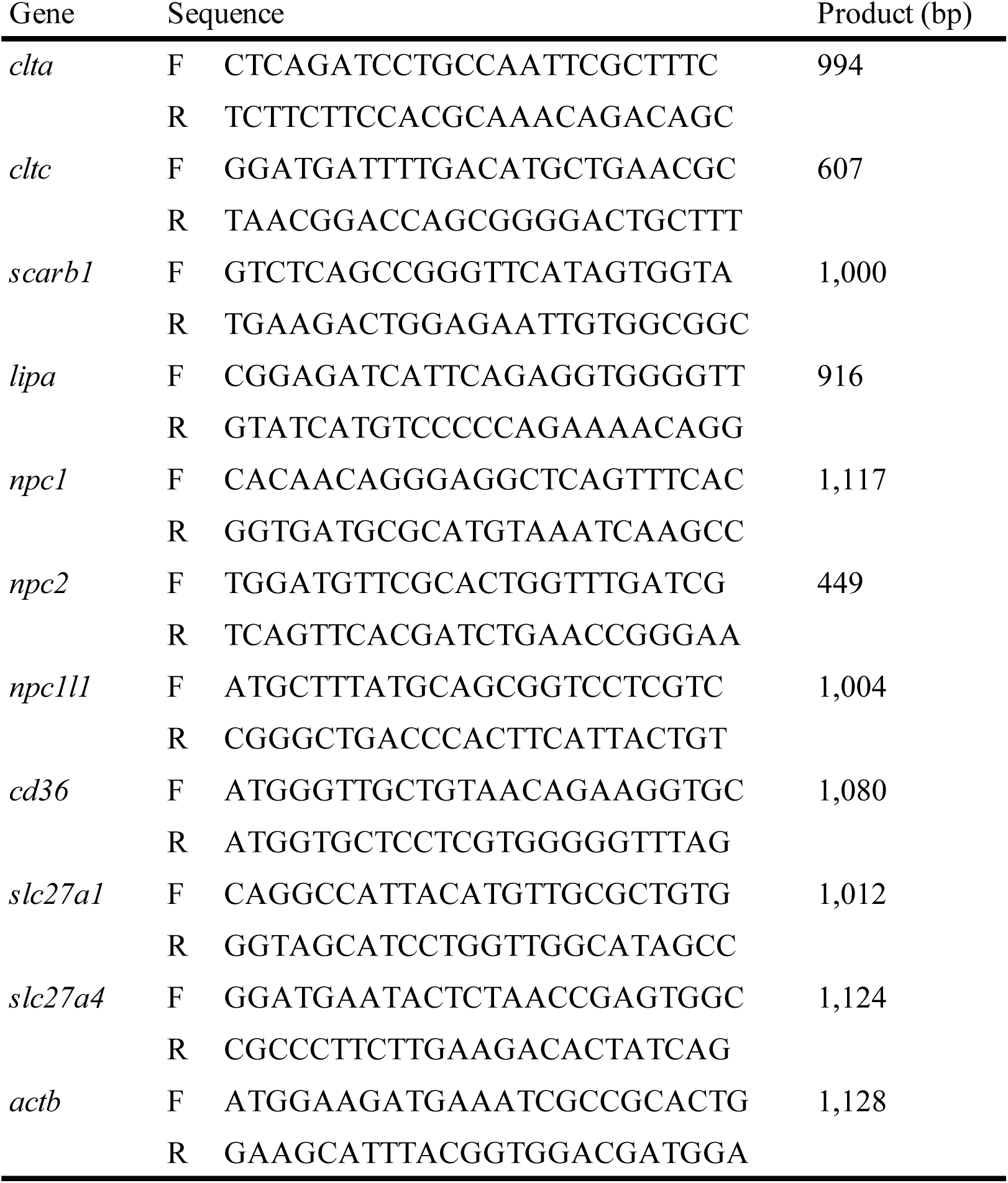
The primers for RT-PCR.

**Table S5.**
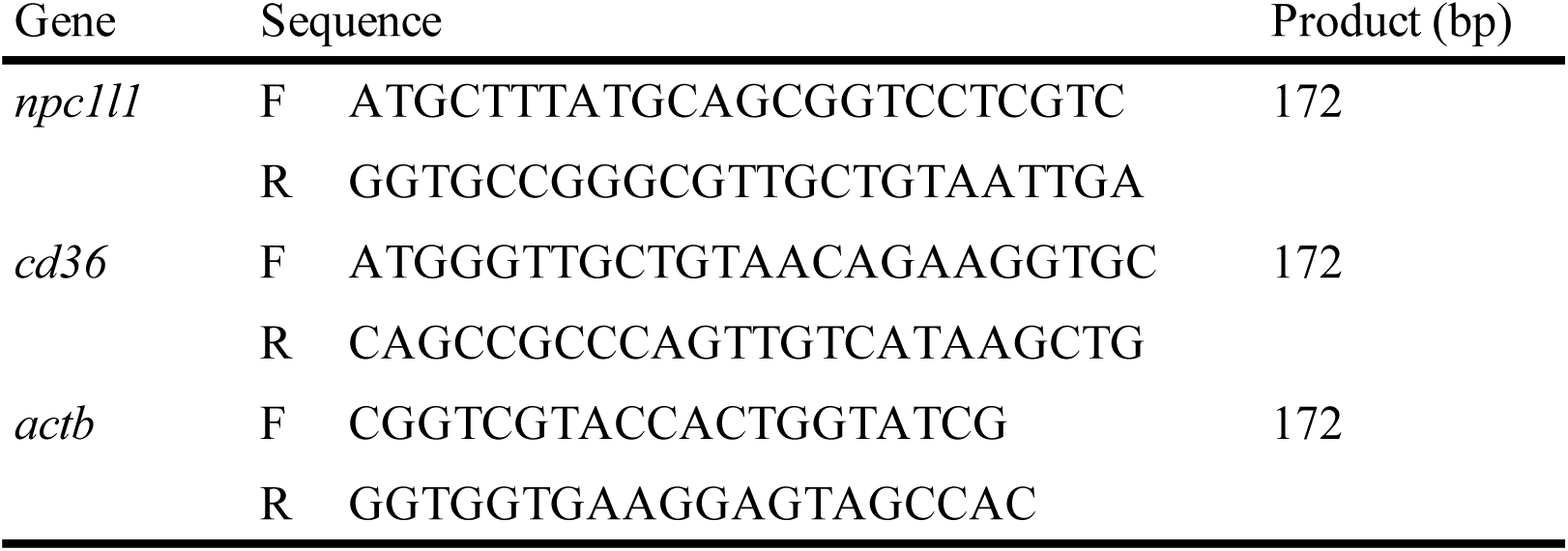
The primers for qPCR.

**Table S6.**
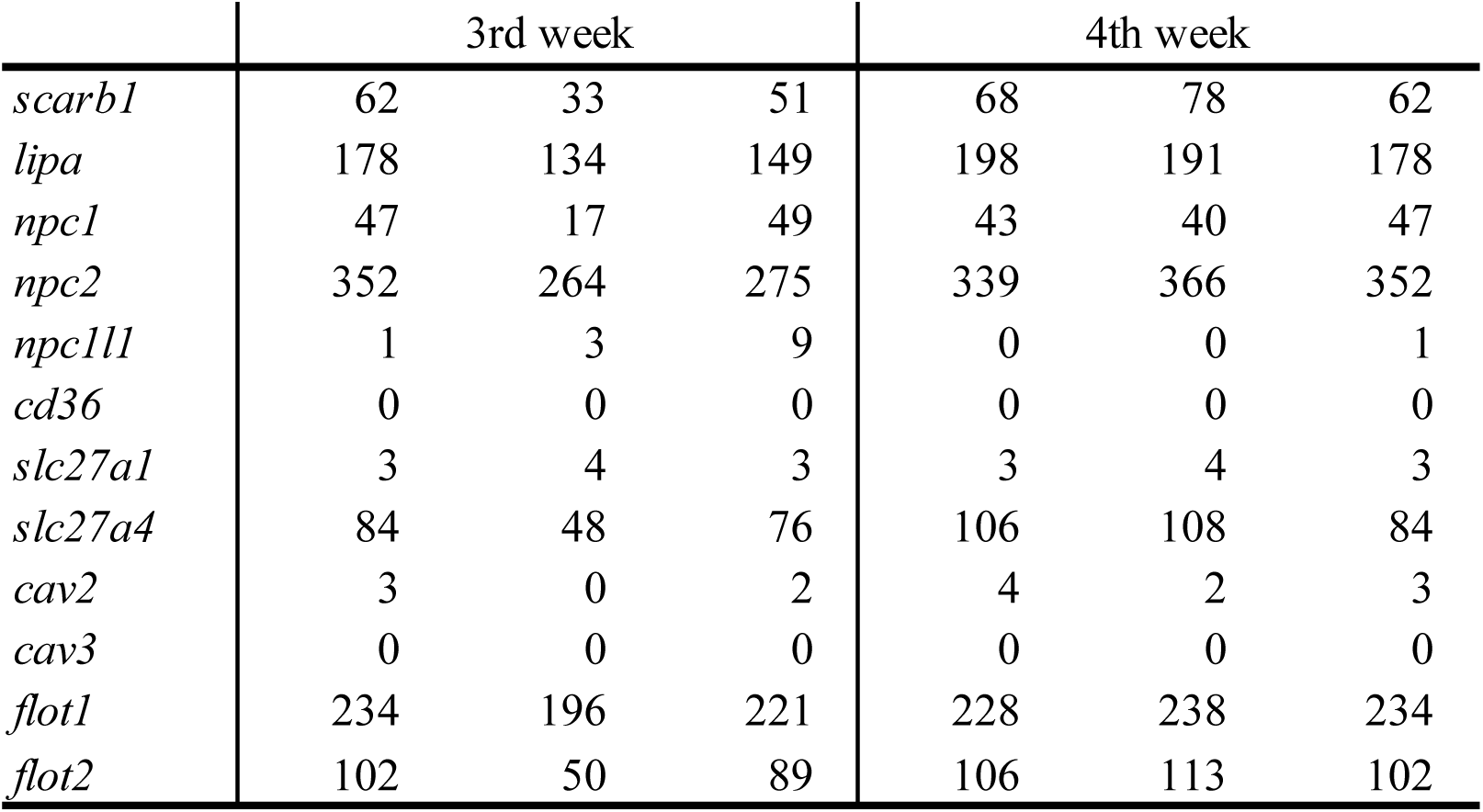
The TPM values based on the open transcritome data.

